# Kronos: a workflow assembler for genome analytics and informatics

**DOI:** 10.1101/040352

**Authors:** M Jafar Taghiyar, Jamie Rosner, Diljot Grewal, Bruno Grande, Rad Aniba, Jasleen Grewal, Paul C Boutros, Ryan D Morin, Ali Bashashati, Sohrab P Shah

**Author notes:** Correspondence &.

## Abstract

**Background:** The field of next generation sequencing informatics has matured to a point where algorithmic advances in sequence alignment and individual feature detection methods have stabilized. Practical and robust implementation of complex analytical workflows (where such tools are structured into ‘best practices’ for automated analysis of NGS datasets) still requires significant programming investment and expertise.

**Results:** We present *Kronos*, a software platform for automating the development and execution of reproducible, auditable and distributable bioinformatics workflows. Kronos obviates the need for explicit coding of workflows by compiling a text configuration file into executable Python applications. The framework of each workflow includes a run manager to execute the encoded workflows locally (or on a cluster or cloud), parallelize tasks, and log all runtime events. Resulting workflows are highly modular and configurable by construction, facilitating flexible and extensible meta-applications which can be modified easily through configuration file editing. The workflows are fully encoded for ease of distribution and can be instantiated on external systems, promoting and facilitating reproducible research and comparative analyses. We introduce a framework for building Kronos components which function as shareable, modular nodes in Kronos workflows.

**Conclusion:** The Kronos platform provides a standard framework for developers to implement custom tools, reuse existing tools, and contribute to the community at large. Kronos is shipped with both Docker and Amazon AWS machine images. It is free, open source and available through PyPI (Python Package Index) and https://github.com/jtaghiyar/kronos.

## Background

The emergence of next generation sequencing (NGS) technology has created unprecedented opportunities to identify and study the impact of genomic aberrations on genome-wide scales. Data generation technology for NGS is stabilizing and exponential declines in cost have made sequencing accessible to most research and clinical groups. Alongside progress in data generation capacity, a myriad of analytical approaches and software tools have been developed to identify and interpret relevant biological features. These include computational methods for raw data pre-processing, sequence alignment and assembly, variant identification, and variant annotation. However, major challenges are induced by rapid development and improvement of analytical methods. This makes construction of analytical workflows a near dynamic process, creating a roadblock to seamless implementation of linked processes that navigate from raw input to annotated variants. Most workflow solutions are bespoke, inflexible, and require considerable programming and software development for their implementation. Consequently, the field currently lacks software platforms that facilitate the creation, updating, and distribution of workflows for advanced and reproducible data analysis by clinical and research labs. Robust analysis of large sets of sequencing data therefore remains labor intensive, costly, and requires considerable analytical expertise. As best practices (*e.g.*, [1]) remain a moving target, software systems that can rapidly adapt to new (and optimal) solutions for domain-specific problems are necessary to facilitate high-throughput comparisons.

Several tools and frameworks for NGS data analysis and workflow management have been developed to address these needs. Galaxy [2], is an open, web-based platform to perform, reproduce and share analyses. Using the Galaxy user interface, users can build analysis workflows from a collection of tools available through the Galaxy toolshed (https://toolshed.g2.bx.psu.edu). The Taverna suite [3] allows the execution of workflows that typically mix web services and local tools. Tight integration with myExperiment [4] gives Taverna access to a network of shared workflows, including NGS data processing. The above tools are mainly aimed at users with minimal programming experience. In addition, Galaxy imposes considerable preparation and installation overhead, lacks explicit representation of workflows (such as in XML format) [5] and imposes some restrictions (such as in file management). Taverna mainly provides a way to run web services and lacks support for scheduling in high performance computing clusters [5].

Due to these limitations, experienced bioinformaticians commonly work at a lower programming level and write their own workflows in scripting languages such as Bash, Perl, or Python [6]. A number of lightweight workflow management tools have been specifically developed to simplify scripting for these target users, including Ruffus [7], Bpipe [8], and Snakemake [9]. While these workflow management tools reduce development overhead, users still need to write a substantial amount of code to create their own workflows, maintain the existing ones, replace subsets of workflows with new ones, and run subsets of existing workflows.

To further facilitate the process of creating workflows by power users, Omics-Pipe proposed a framework to automate best practice multi-omics data analysis workflows based on Ruffus [10]. It offers several pre-existing workflows and reduces the development overhead for tracking the run of each workflow and logging the progress of each analysis step. However, it is remains cumbersome to create a custom workflow with Omics-pipe as users need to manually write a Python script for the new workflow by copying/pasting a specific header to the script and writing the analyses functions using Ruffus decorators. The same applies when adding or removing an analysis step to an existing workflow.

We introduce a highly flexible open-source Python-based software tool (Kronos), that significantly reduces programming overhead for workflow development. Kronos has a built-in run manager that parallelizes subsets of the workflow specified by the user, logs the runtime events (provides full analysis chain of custody), and relaunches a workflow from where it left off. It can also execute the resulting workflow locally, on a compute cluster or cloud. The workflows generated by this tool are highly modular and flexible. Changing a workflow by adding, removing or replacing analysis modules (referred to as *components*), or altering the analysis parameters can be easily achieved by reconfiguring the configuration file (without having to manually modify the source code of the workflow). The configuration files and *components* are shareable; therefore, users can readily regenerate a workflow elsewhere, facilitating reproduciblity. In addition, Kronos has a framework for creating new *components* that can be easily shared and reused by collaborators or others in the bioinformatics community. Kronos is shipped with Docker and Amazon Machine images to further facilitate its use locally, on high performance computing clusters and in the cloud infrastructures. Instantiated workflows and *components* for the analysis of single human genomes and cancer tumour-normal pairs following best analysis practices accompany Kronos and are freely available.

## Results

Kronos transforms a set of existing *components* (i.e., analysis modules; described later) along with a configuration file into a modular workflow without having to write code. It also provides a functionality to create *component* templates which greatly facilitates developing *components* by experienced bioinformaticians.

As shown in Figure 1, users can conveniently create a workflow by following three steps listed below (referred to as Steps 1, 2 and 3 in the remainder of this paper). Section 2 of Additional file 1 provides an example of how to make a variant calling workflow.

**Figure 1.**
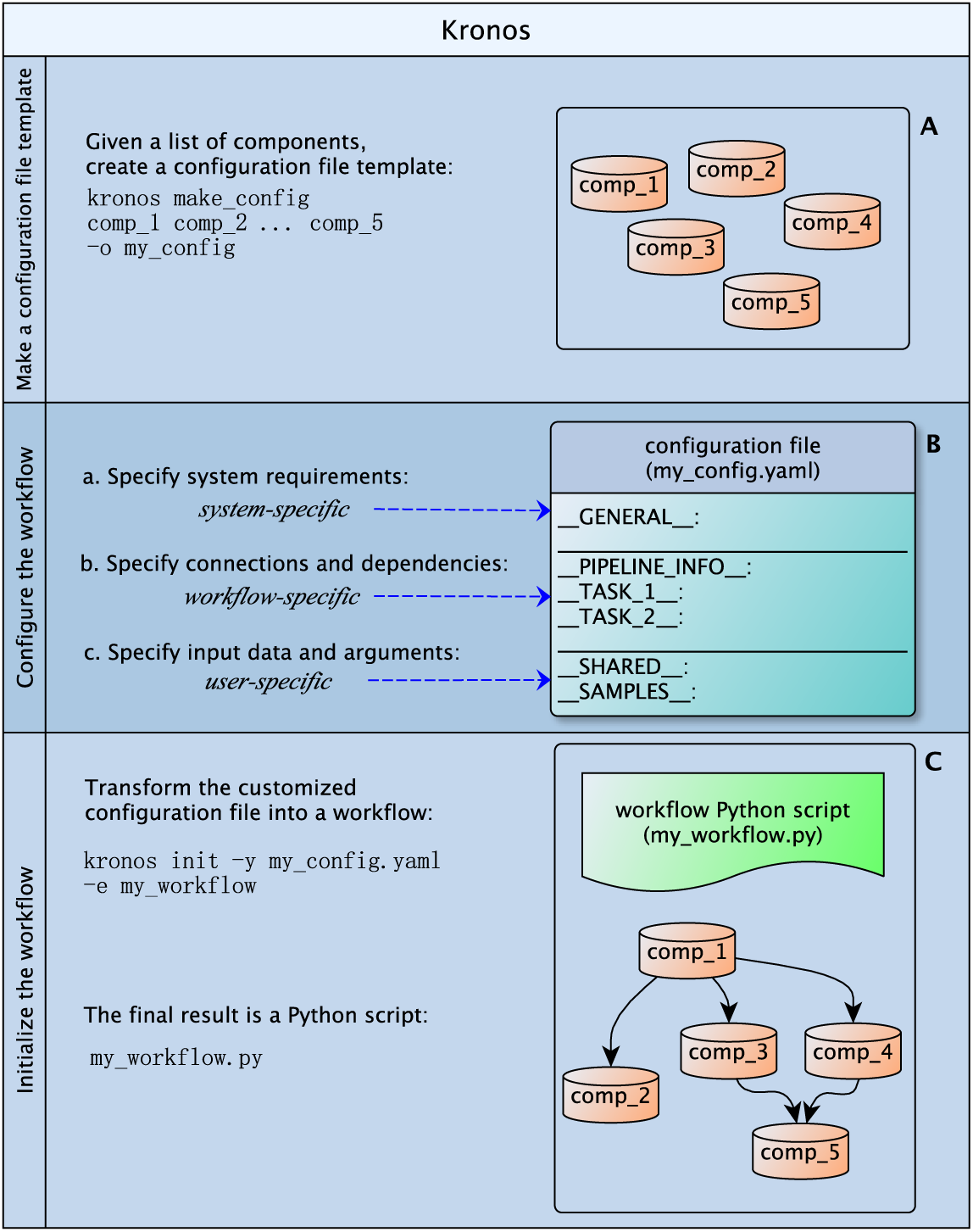
Make a workflow. Making a new workflow with Kronos includes three steps: i) make a configuration file template: given a set of existing *components*, users can generate this file by running the command `make_config`; ii) configure the workflow: users can specify the desirable flow of their workflow using the connections and dependencies, customize output directory names, and specify input arguments and data to the required fields in the configuration file template; iii) initialize the workflow: this is achieved by running the command `init` on the configuration file which transforms the YAML file into the Python workflow script.

- Step 1. Given a set of existing *components*, create a configuration file template by running the following Kronos command:

kronos make_config

[ list of components ] –o <output name>

where [list of components] refers to the *component* names that we aim at using in our workflow.
- Step 2. In the configuration file template, specify the order by which the *components* in the workflow should be run. This does not require programming skills and is merely text-based.
- Step 3. Create the workflow by running the following Kronos command with the configuration file as its input:

kronos init– y <config file.yaml>

–o <workflow _name>

The output is an executable Python script that uses the built-in run manager of Kronos. The run manager provides scalability by enabling users to run the workflow on a single machine, on a cluster of computing nodes or in the cloud. In fact, each *component* in the workflow can individually be run either locally, on a cluster. In addition, it allows users to independently set native specifications such as free memory, maximum memory or the number of CPU’s, for each task.

The run manager also provides the following features for the resulting workflow:

- generates a unique run-ID for each run
- re-runs the workflow from where it left off using the run-ID
- runs intermediate workflows in parallel
- limits the number of concurrent jobs and workflows as desired
- creates a detailed log file for each run tagged with the run-ID

### Kronos *components*

In order for different software tools, referred to as *seeds*, to be used as input to the `make_config` command (Step 1), users need to wrap them with a number of particular files. We call a wrapped *seed* a *component*. A *seed* can be as simple as a command copying a file or it can be a more complicated tool such as a single nucleotide variant (SNV) caller.

Regardless of how complicated a *seed* is, its corresponding *component* has a standard directory structure composed of specific wrappers and sub-directories. The wrappers are independent of the programming language used for developing the *seed* and essentially all tools can be wrapped as *components*. In addition, Kronos provides a functionality (through `make_component` command) to create a *component* template which helps developing a new *component* in a few minutes (Figure 2) provided that the *seed* exists. This process is straight-forward and requires minimal programming, yet it provides a powerful framework for experienced programmers to fully customize their workflows. Section 1 of Additional file 1 provides an example creating a *component*.

**Figure 2.**
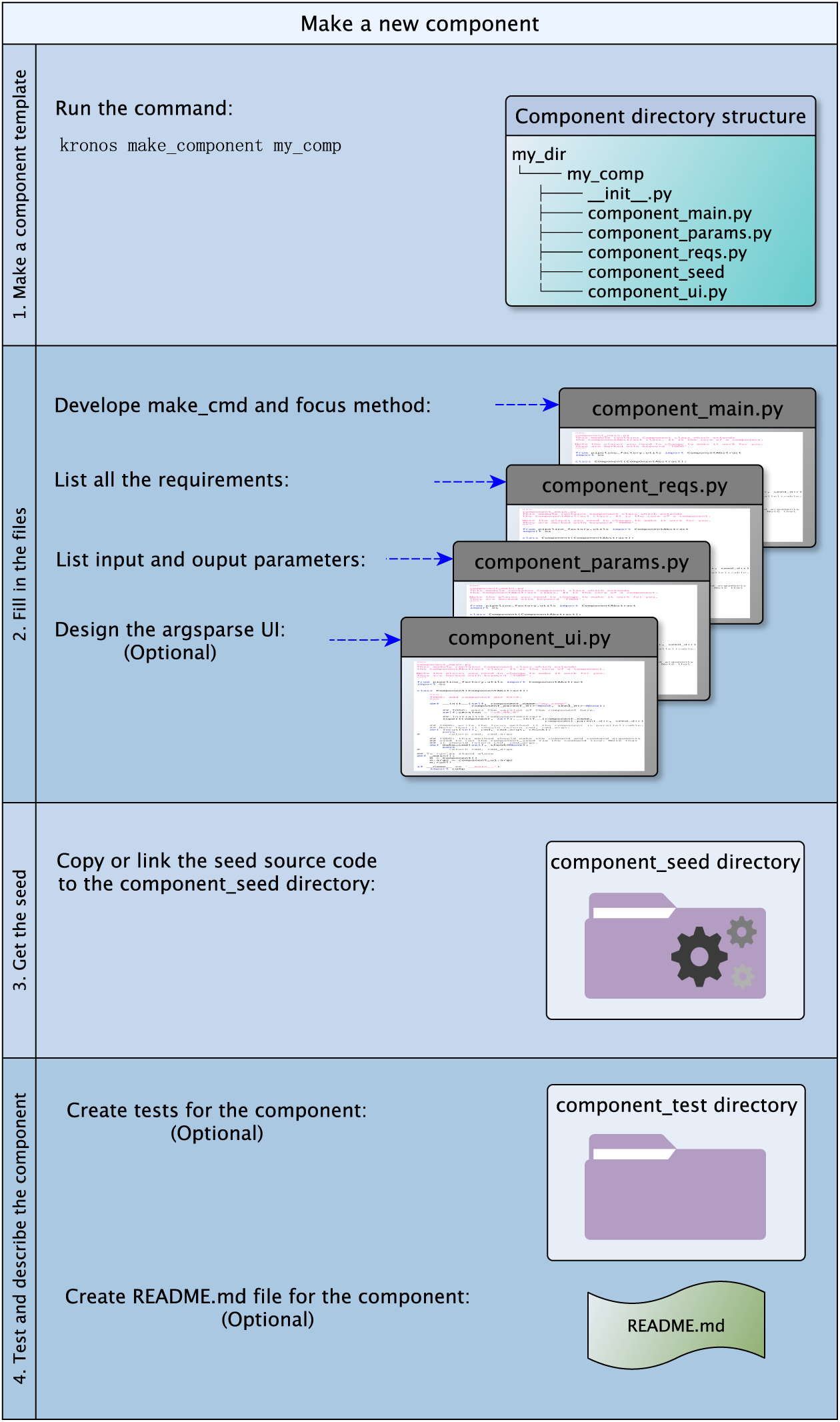
Make a component. Making a new *component* for Kronos includes the following steps: i) make a new *component* template by running the command `make_component`; ii) fill in the resulting template accordingly; iii) copy or link the source code of the *seed* used in the *component*; iv) optionally create `README.md` and tests for the *component*.

### Kronos configuration file

The `make_config` command generates a configuration file template. It is a text file formatted as YAML and contains all the parameters of the input *components* which are mostly pre-filled with default values. For each input *component*, there is a corresponding section with a unique name in the configuration file called *task*. Users should use these sections to specify the order by which each *task* in the workflow should be run (Step 2 of creating a workflow). This can be done by a simple convention called *IO-connection*. An *IO-connection* is basically a pair of values comprising of a *task* name and one of its parameters. It determines which *task* should be followed by the current *task* and is specified as an argument to one of the parameters of the current *task*. For example, in the following configuration file, `(’_TASK_1_’, ‘out_file’)` is an *IO-connection* which makes `_TASK_2_` to follow `_TASK_1_`, *i.e*. the input to the parameter `in_file` of `_TASK_2_` comes from the parameter `out_file` of `_TASK_1_`.

_TASK_1_:

out _file: ‘foo.txt’

_TASK_2_:

in file: (’ _TASK_1_’, ‘out file’)

A configuration file has the following blocks (see Additional file 1: Figure S1):

- system-specific which captures the system dependant requirements of the workflow (such as the paths to the local installations) and includes the `GENERAL` and `PIPELINE_INFO` sections.
- user-specific which contains the input files and arguments and includes the `SHARED` and `SAMPLES` sections.
- workflow-specific which defines the connection between the *components* in the workflow. *Task* sections related to each *component* are in this group.

This design has the following advantages: i) if users want to run the same workflow for various sets of input files and arguments, they would only need to update the user-specific sections. This prevents inadvertent changes in the flow of the workflow when changing the inputs; and ii) the segregation of system-specific information from the rest of the sections enables users to run a workflow practically anywhere. In other words, by simply updating the system-specific sections with proper values, the requirements of the workflow can be observed on any machine or cluster.

### Kronos workflows

Each workflow made by Kronos is a directed acyclic graph (DAG) of *components* where every node in the graph corresponds to a *task* section in the configuration file. *Task* sections can independently be added, removed or replaced in the configuration file (Figure 3). Therefore, to add, remove or replace a *component* in the workflow or equivalently a node in the DAG, users simply need to change the corresponding *task* section in the configuration file and run the command in Step 3. As a result, the workflows are highly modular and maintaining them is as easy as updating the configuration file without having to rewrite the workflow. Finally, a workflow can be run by simply running the Python workflow script using the command-line as depicted in Figure 4.

**Figure 3.**
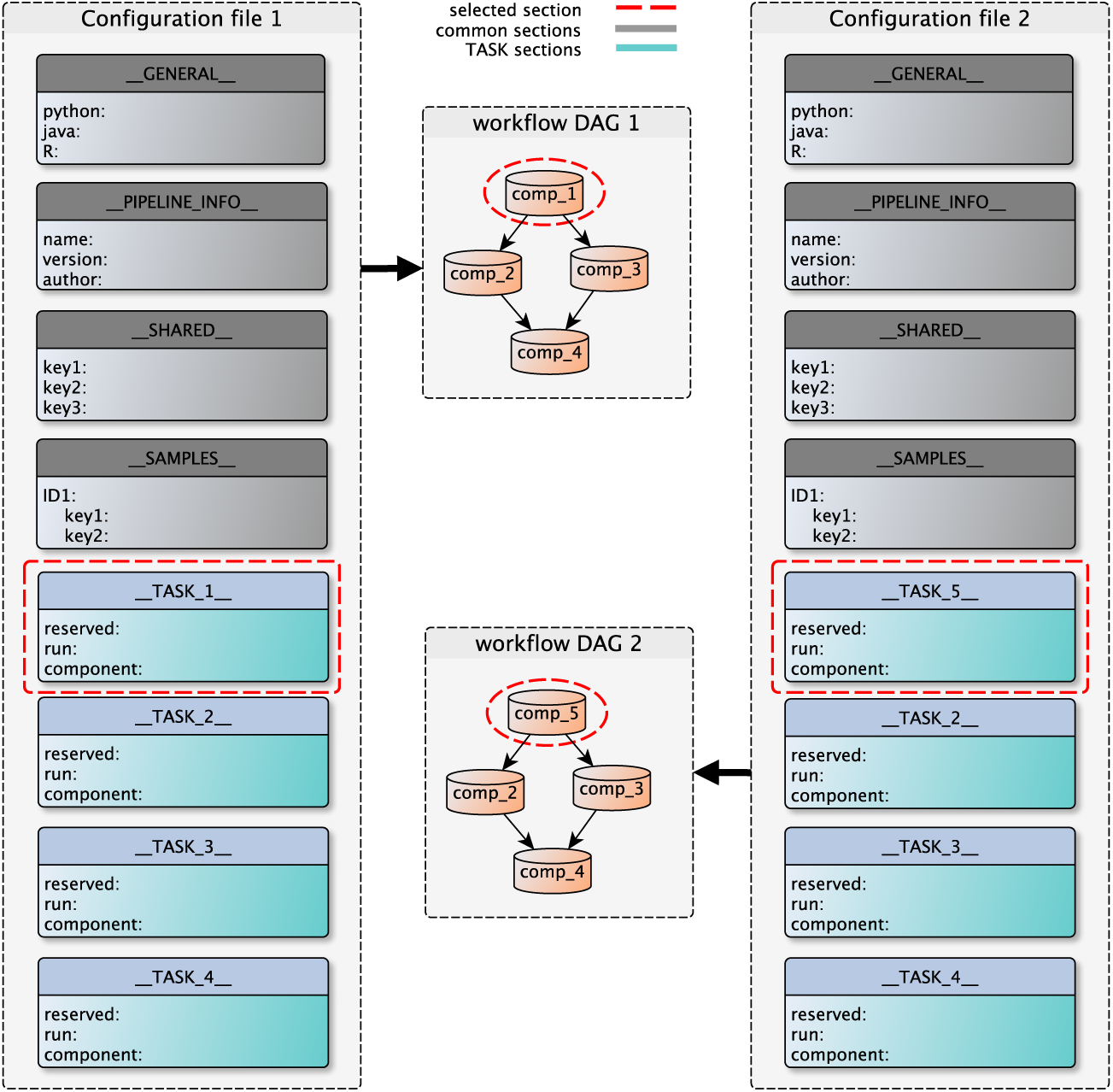
Replace a component in a workflow. The configuration file has different sections as shown in the figure. These sections are: GENERAL, PIPELINE INFO, SHARED, SAMPLES, and TASKs. The modular organization of the configuration file allows for easy customization of workflows which can serve different purposes such as tool comparison. Adding, removing or replacing nodes in the DAG of the workflows can be easily done by adding, removing or replacing the corresponding TASK sections in the configuration file. For instance, to go from workflow `DAG1` to workflow `DAG2`, *i.e*. to replace `comp_1` (*e.g.*, variant caller 1) in the first workflow by comp_5 (*e.g.*, variant caller 2) in the second, the user only needs to replace `TASK_1` section by `TASK_5` section in the configuration file and perform Step 3.

**Figure 4.**
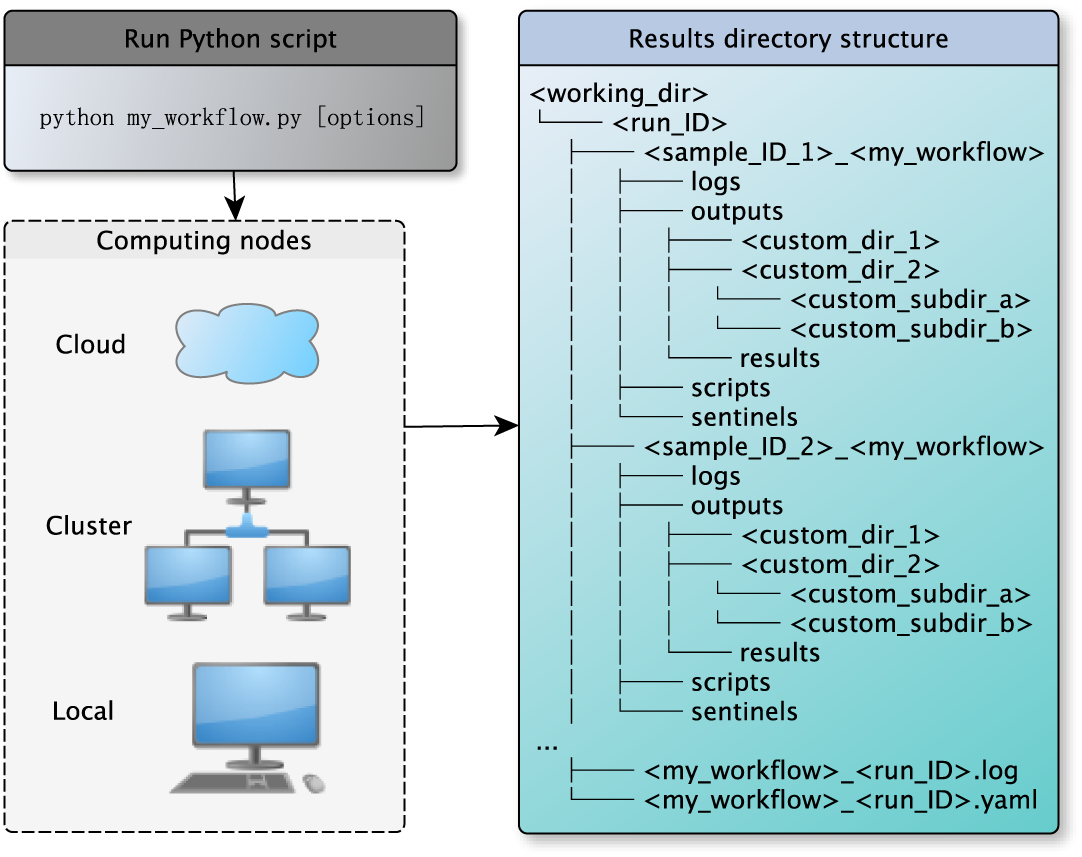
Run a workflow. Workflows generated by Kronos are ready to run locally, on a cluster of computing nodes, and in the cloud. To run a workflow, users only need to run the Python workflow script. Each run of a workflow generates a specific directory structure tagged with a run-ID. When running a workflow for multiple samples, a separate directory is made for each sample to make it convenient to locate the results corresponding to each sample. This figure shows the tree structure of the resulting directory. There are four sub-directories that are always generated for each sample: i) logs: to store the log files; ii) outputs: to store all the output files generated by all the *components* in the workflow; iii) scripts: to store the scripts automatically generated by Kronos to run each *component* in the workflow; iv) sentinels: to store sentinel files used by Kronos to pick up the workflow from where it left off in a previous run.

### Kronos features and benefits

Full details of how to use each of the following features can be found in the software documentation.

#### Parameter sweeping

It is sometimes desired to run a particular tool or algorithm with various sets of parameters in order to select the parameter set with superior performance for a given problem. For example, a user may want to find the proper model parameters (such as mapping quality and base quality thresholds) for a variant calling tool to accurately detect single nucleotide variants. Kronos provides a mechanism for this purpose where users can specify all different sets of input arguments (or parameters) in the `SAMPLES` section of the configuration file. In this case, running Step 3 creates a number of intermediate workflows, each for one set of input arguments, along with the main workflow. When running the main workflow, Kronos run manager automatically runs the intermediate workflows in parallel, each on one set of the input arguments. We have provided a variant calling workflow with parameter sweeping functionality in Section 3 of Additional file 1 to demonstrate this feature.

#### Tool comparison

In bioinformatics, it is often required to compare the performance of two or more algorithms or compare a new analysis tool to the existing ones to select the best that fits the particular goals of a project. For example, it is often helpful to evaluate the performance of different variant calling algorithms [11]. The modularity of the workflows generated by Kronos facilitates the comparison of different algorithms and tools. For this purpose, as shown in Figure 3, the user can simply replace a *task* section corresponding to an analysis tool with another *task* section corresponding to another similar tool and run Step 3.

#### Reproducibility

The configuration file and *components* of a workflow made by Kronos are portable. Therefore, users can readily re-create the same workflow by only replacing the system-specific section of the configuration file and running the command in Step 3. To show this functionality, we have included an example of a workflow that performs somatic variant calling on whole genome data of a breast cancer case using Strelka algorithm [12] and generates a number of plots based on Strelka calls (Figure 5). Detailed step-by-step instructions to reproduce this figure is in Section 3 of Additional file 1.

**Figure 5.**
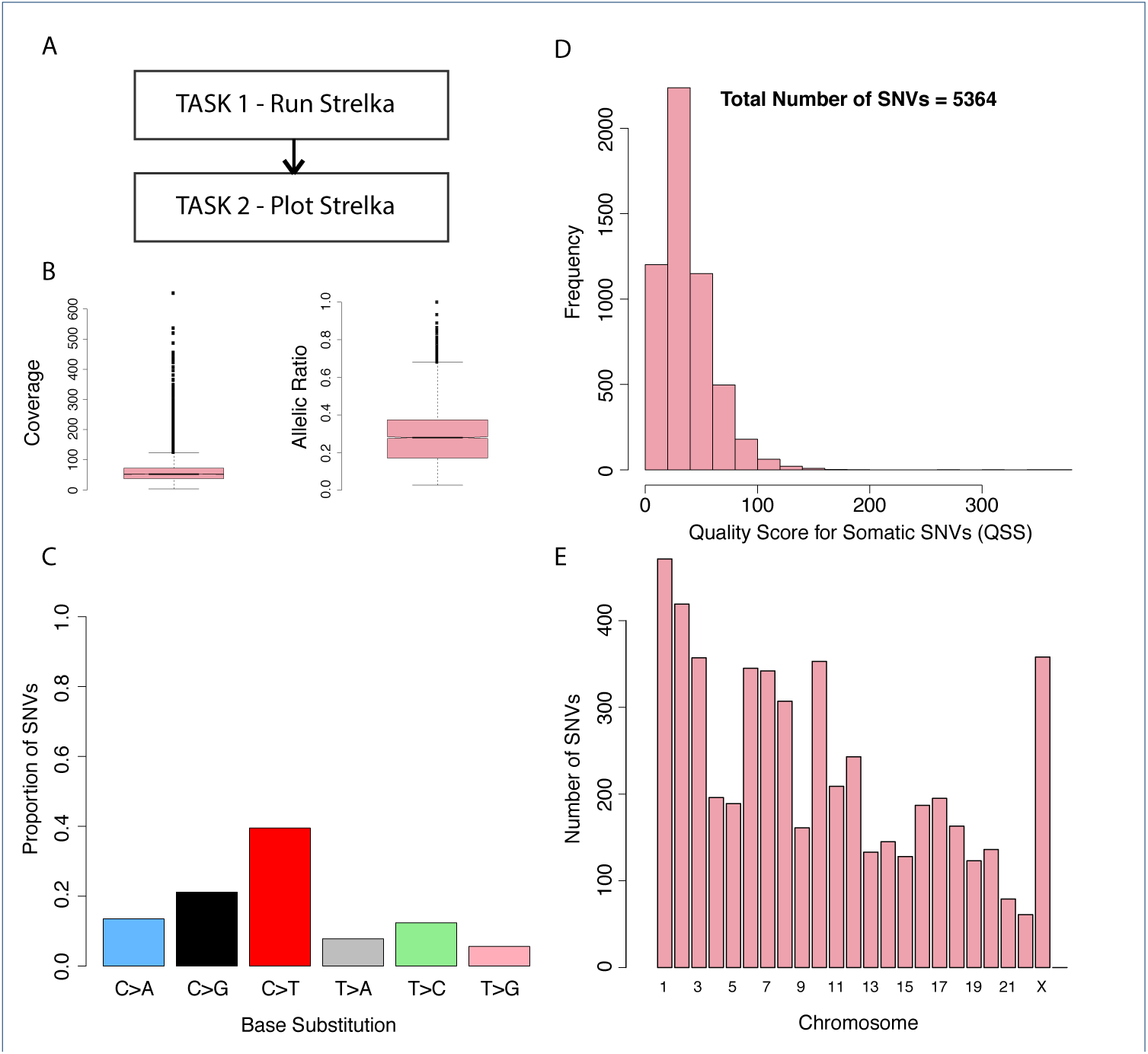
Strelka workflow. Results from the tumour-normal variant calling workflow on whole genome data of a breast cancer case (SA500 - EGA accession number EGAS00001000952). (A) schematic of the workflow which is comprised of two tasks. The plots generated by the workflow is in fact the output of `TASK_2`, (B) box plot of coverage and variant allelic ratios for the SNVs detected by Strelka, (C) base substitution patterns for the somatic SNVs, and (D) total number of SNVs and their histogram based on the quality score (QSS), (E) Distribution of the number of SNVs across different chromosomes.

#### Automatic parallelization and merge

Most of the recent tools developed in bioinformatics field are parallelizable or have the potential to run in parallel. However, majority of these tools are shipped without the built-in functionality and require the users to manually break the analysis into smaller analyses. For example, many variant calling algorithms are capable of running on user-specified coordinates of the genome but are not shipped with parallelization functionality. However, a user can analyze a whole genome sequencing data chunk by chunk in parallel with the caveat of manually scripting the parallelization steps. Due to the cumbersome nature of manual parallelization, many users might avoid running the tools in parallel which considerably increases the runtime of the analysis. To resolve this issue, Kronos automatically parallelizes tasks in the workflow if feasible. Then, it aggregates the outputs of all child tasks and merges them if necessary.

#### Cloud support

The massive scale of genomic data is justifying a move to the cloud for storage and analyses in order to minimize cost and handle the ebb and flow of computational demands. Kronos’ flexibility addresses the emerging need for rapid deployment of analysis workflows in the cloud. Several command-line tools exist for managing fleets of compute nodes on cloud platforms such as Amazon Web Services (AWS), including StarCluster, CfnCluster and Elasticluster. Similarly, the Galaxy project provides CloudMan, a graphical interface for launching so-called “cloud clusters”. Kronos is scheduler-agnostic; therefore, developers can leverage its powerful features in combination with any of these tools. A guide on creation and management of a cloud cluster using the StarCluster software and deployment of Kronos is provided in the online documentation and an Amazon Machine Image (AMI) is provided for convenience.

#### Controlled pause/resume by breakpoints

When running a workflow, certain blocks of the workflow may need to run multiple times, for example to tune a particular parameter of a *component* or to inspect the results of the previous *tasks* in the workflow before the next *tasks* are triggered. Analogous to the debuggers, Kronos provides users with breakpoints to perform a controlled pause/resume action.

In addition, with the breakpoint mechanism, users can break the flow of a workflow into several sub-workflows and run each part on a different machine or cluster. In other words, once a breakpoint happens, *i.e*. one sub-workflow is complete, the main workflow can be transferred to a different machine and it will pick up running from where it left off on the previous machine provided that all the intermediate files are present. For example, a workflow can contain a *component* as its last step that loads the final results to a local database which can be reached only from a specific IP or machine. In this case, the user can run the workflow on a powerful computing node or a cluster with a breakpoint set for the *component* prior to the last *component*, *i.e*. database loader in this example. Once the breakpoint is applied, the user can resume the workflow on the other machine, so that the results can be loaded to the local database.

#### Forced dependency

Often in a workflow, a *task* requires the output of the previous one. As explained earlier, Kronos handles this explicit dependency by *IO-connection*. However, sometimes a *task* might need to wait for one or more other steps in the workflow to finish although there are no explicit *IO-connections* between them. For example, when two tasks intend to write results in the same file, one needs to make sure that both *tasks* do not run at the same time. Another example would be a variant calling algorithm (*e.g.*, GATK) which accepts a bam file as input. However, it also expects the index of the bam file to be present in the same directory as the bam file. If the index is created in one of the previous *tasks* in the workflow, then the current *task* that needs the bam file and its index, would depend implicitly on the other *task* that creates the index file. In this case, a mechanism is required to force the variant calling *task* to wait until the index file is ready. Kronos provides forced dependency feature to overcome this problem (see Additional file 1: Figure S2).

#### Results directory customization

It is desirable to have full control of the structure of the results directory when running a workflow. With Kronos, users can readily determine the structure of the results directory in the configuration file. This provides an easy file management for the users. Figure 4 shows an example of the tree structure of the results directory generated for a workflow.

#### Boilerplates

Users can use this feature to insert a command or a script into the begining of the command used to run a *task* in a workflow. This is particularly useful for setting up the environments using the Environment Modules package [13]. It also provides a means to run preprocessing steps for a specific *task* prior to running the *task* itself.

#### Keywords

There are several specific keywords that users can use in the configuration file which will be automatically replaced by proper values in runtime. This enables users to customize the paths and file names based on the workflow-specific values in runtime such as run-ID, workflow name or sample ID.

## Workflows

We have developed a number of standard genome analysis workflows using Kronos. These workflows utilize many of the Kronos features introduced earlier and are publicly available.

### Alignment workflow

This workflow accepts paired-end FASTQ files as input and aligns them using the Burrows-Wheeler aligner [14]. It also sorts the aligned bam file, flags the duplicates, indexes the file and generates statistics for the final bam file.

### Germline variant calling workflow

This workflow is an implementation of the best practices guide established by the Broad Institute [1] applied to variant discovery using haplotypecaller. In short, it runs the Bowtie2 aligner, creates targets using GATK RealignerTargetCreator, and calls SNVs and indels using GATK.

### Copy number estimation workflow

HMMcopy is a suite of tools for copy number estimation of whole genome sequencing data [15]. This workflow takes a bam file as an input and estimates the copy number with GC and mappability correction using HMMCopy. It also segments and classifies the copy number profiles with a robust Hidden Markov Model.

### Somatic variant calling workflow

This workflow takes a pair of tumour/normal bam files as inputs and detects the somatic SNVs and indels using Strelka algorithm [12], annotates the resulting VCF files using SnpEff [16], and flags the variants observed in 1000 genomes and dbSNP databases.

### RNA-seq analysis workflow

This workflow aligns RNA-seq FASTQ files using STAR aligner [17] followed by Cufflinks which assembles transcriptomes from RNA-Seq data and quantifies their expression [18].

## Conclusions

A foundation for rapid and reliable implementation of genomic analysis workflows is an essential need as a myriad of potential applications of genomics (ranging from personalized cancer therapies to monitoring the evolution and spread of infectious diseases) are projected to produce massive amount of genomic data in the next few years. We have developed Kronos to address this need and standardize reproducibility of genome analysis tasks. Kronos minimizes the cumbersome process of writing code for a workflow by transforming a YAML configuration file into a Python script and manages workflow execution locally, on a cluster or in the cloud. Given a set of pre-made *components*, constructing a workflow by Kronos does not need programming skills as the user only needs to fill out specific sections of a YAML configuration file. Making *components* still requires programming. However, component development time and effort is minimal given the design structure of Kronos’ *components*.

In addition, Kronos provides a functionality to create a *component* template that can be used to wrap an existing software (*seed*) with minimal coding intervention. This provides a powerful and highly flexible framework for bioinformaticians to fully customize their workflows. A number of standard genomic analysis workflows along with their building *components* that have been made by Kronos accompany this software and are available to public. This resource not only provides users with a suite of workflows for standard genomics analysis, but also introduces a framework for power users to develop custom *components*, reuse or tweak existing ones, and finally share them with their collaborators and community at large. Kronos has been developed for genomics applications but it can be readily utilized in other scientific and non-scientific fields.

In conclusion, this work provides a framework towards rapid integration of new (and optimal) genomic analysis advances in high-throughput studies. The flexibility, customization, and modularity of Kronos make it an attractive system to use in any high-throughput genomics analysis endeavour. We expect Kronos will provide a foundational platform to accelerate towards the need to standardize and distribute NGS workflows in both clinical and research applications.

## Availability and requirements

Kronos is a free and open-source Python package available through PyPI (Python Package Index) under the MIT license. Documentation can be found at https://readthedocs.org/projects/kronos/ and the workflows and their components accompanying this paper are available at https://github.com/MO-BCCRC?tab=repositories.

## Competing interests

The authors declare that they have no competing interests.

## Author’s contributions

J.T. developed the software, wrote the documentation and contributed to manuscript writing. J.R. assisted in developing part of the logger and a few of the helper functions for the software, testing software features and providing feedback on the manuscript. D.G. developed a number of pipelines accompanying the manuscript, tested the software and provided feedback on the software features and manuscript. B.G. deployed and tested Kronos in the cloud, wrote the documentation for cloud deployment, tested software and provided feedback on the software features and manuscript. R.A. developed the germline variant calling workflow and provided feedback on the manuscript. J.G. tested and provided feedback on the software. P.B. provided feedback on the manuscript. R.M. provided resources and supervised testing Kronos in the cloud, and provided feedback on the manuscript. A.B. contributed to the design and development of the software. A.B. and S.S. co-supervised, provided intellectual contributions to the work and contributed to manuscript writing. A.B. and S.S. are joint senior authors.

## Acknowledgements

The authors would like to thank Shadrielle Melijah G. Espiritu and Andre Masella for their feedback on the manuscript/software. This project has been supported by funding from Genome Canada/Genome British Columbia (grant No. 173CIC), Natural Science & Engineering Research Council of Canada (grant No. RGPGR 488167-2013), and Terry Fox Research Institute - Program Project Grants (grant No. 1021).

## Figures

### Additional Files

Additional file 1 — Supplementary information

The supplementary information is in pdf format and expalins a) how to make a *component*, b) how to make a workflow, c) how to run a workflow, and d) Figure S1 and Figure S2.

